# Three-dimensionally preserved ‘Stage IIIb’ fossil down feather supports developmental modularity in feather evolution

**DOI:** 10.1101/2020.08.26.268060

**Authors:** Arindam Roy, Case V. Miller, Michael Pittman, Thomas G. Kaye, Adolf Peretti

## Abstract

We describe a unique three-dimensionally preserved fossil down feather from the Late Cretaceous of Myanmar. It’s morphology is highly congruent with Stage IIIb of the widely accepted Prum and Brush model of feather evolution-development. This makes the new specimen the first evidence of this developmental stage in the fossil record. The Stage IIIb diagnosis is robustly supported by the absence of a central rachis and by its paired barbules emanating from radially positioned barbs that are attached to a short calamus. Prum and Brush’s model hypothesises a bifurcation in the evolution-development pathway at Stage III. Stage IIIa involves rachis development and branching into barbs. Stage IIIb involves branching of the barbs from the calamus and then further branching of the barbules from the barbs. These two pathways then converge into Stage IIIa+b where feathers produce a rachis, barbs and barbules in nested order, finally leading to Stage IV. Evolution-development studies on the morphogenesis of feathers have unequivocally shown that such feather branching can be controlled by BMP, Noggin, Shh and several other proteins. Therefore, molecular crosstalk can convert a barb into a rachis and vice versa. The topology of this down feather, consistent with specific patterns of modular protein-protein signalling already observed, provides the first definitive evidence that such signalling was responsible for the evolution of a diverse inventory of feather morphologies in non-avialan dinosaurs and early birds since the middle Jurassic.

## Introduction

Late Cretaceous amber from Myanmar (or Burmese amber) provides rare insights into ancient forest ecosystems (see *SI Appendix*). Three-dimensionally preserved fossil feathers recovered from Burmese amber have transformed the study of feather evolution (1). Here we describe a Burmese amber specimen GRS Ref. 32865 containing a distinctive three-dimensionally preserved fossil down feather. We show that this fossil down feather has a unique combination of previously undescribed morphological characteristics (see *SI Appendix*). This allows us to diagnose the feather as the first direct fossil evidence of Stage IIIb in the developmental sequence of feather evolution proposed by Prum and Brush (2).

## Results and Discussion

GRS Ref. 32865 **(Fig.1 A,B)** was diagnosed as a Stage IIIb down feather based on (i) observed feather morphology (absence of a central rachis; paired barbules emanating from radially-positioned barbs that are attached to a short calamus) and (ii) currently available evidence on molecular signalling (protein-crosstalk) occurring within developing feathers **(Fig 2)**. Despite showing a certain degree of morphological disparity, vertebrate integumentary appendages possess widely shared developmental pathways (Wnt, Eda-Edar, BMP’s and Shh) as exemplified in shark denticles, bony fish scales, mammalian hair, reptilian epidermal scales and avian feathers (3). Several attempts have been made to explain the morphogenetic transition of scales and scutes into highly branched feathers (3). Evolution-development experiments show that all integumentary structures originate from embryonic structures in the dermis called placodes (4). The outcome in different vertebrate groups (denticles, hair, scales and feathers) is determined during embryonic development by fine-tuned spaciotemporal regulation of homologous signalling in ectodermal-mesenchymal pathways (5). In birds, experiments have shown feather development has to be inhibited for scales to develop. This indicates that avian scales are secondarily derived and not true analogues to reptilian scales, as confirmed by their different molecular profiles (6).

**Figure 1.**
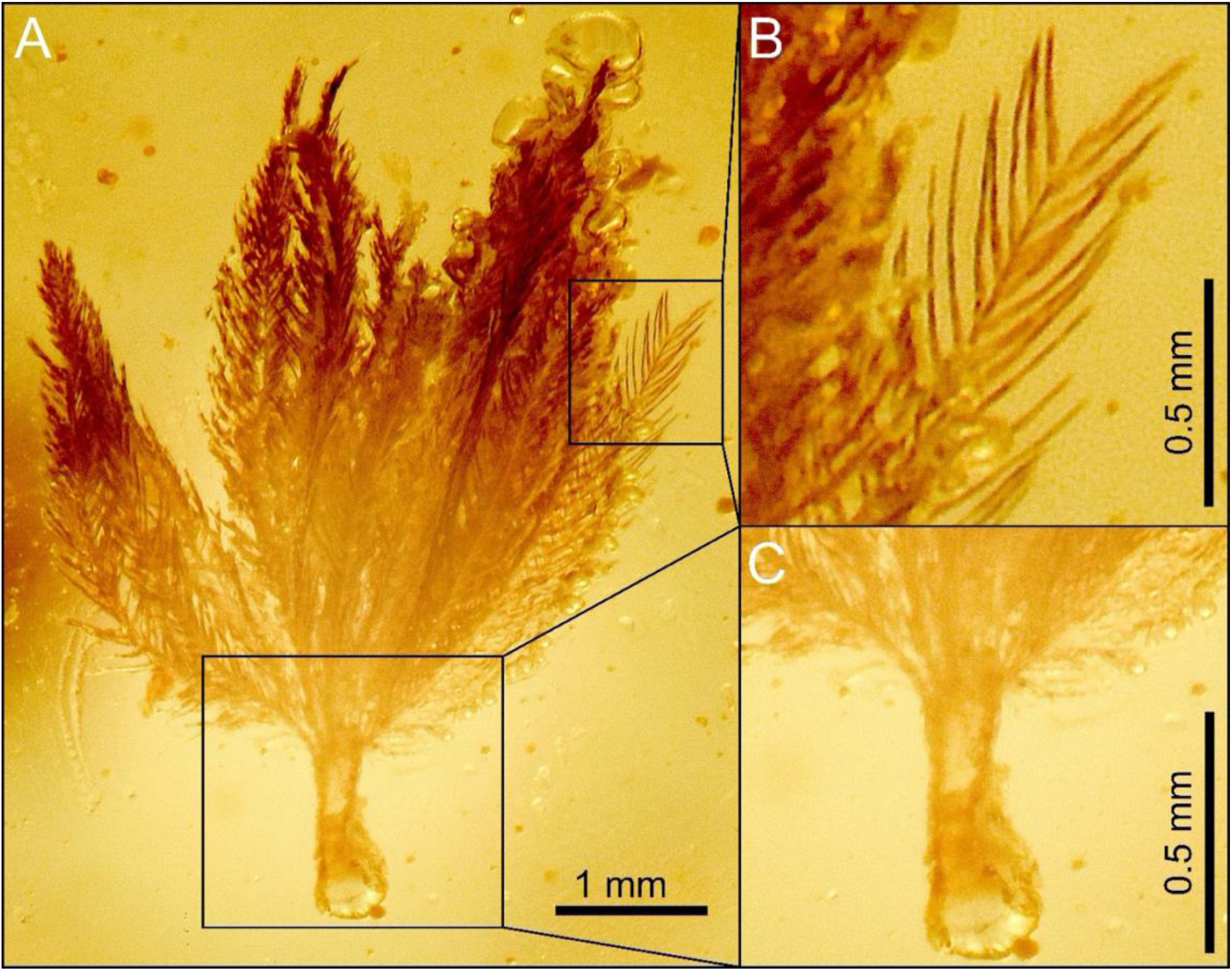
Down feather trapped within amber specimen GRS Ref. 32865. **A**, full view of the feather. **B**, detail of the best-preserved barb and its constituent paired barbules. **C**, detail of the calamus, with proximal bulge.

**Figure 2.**
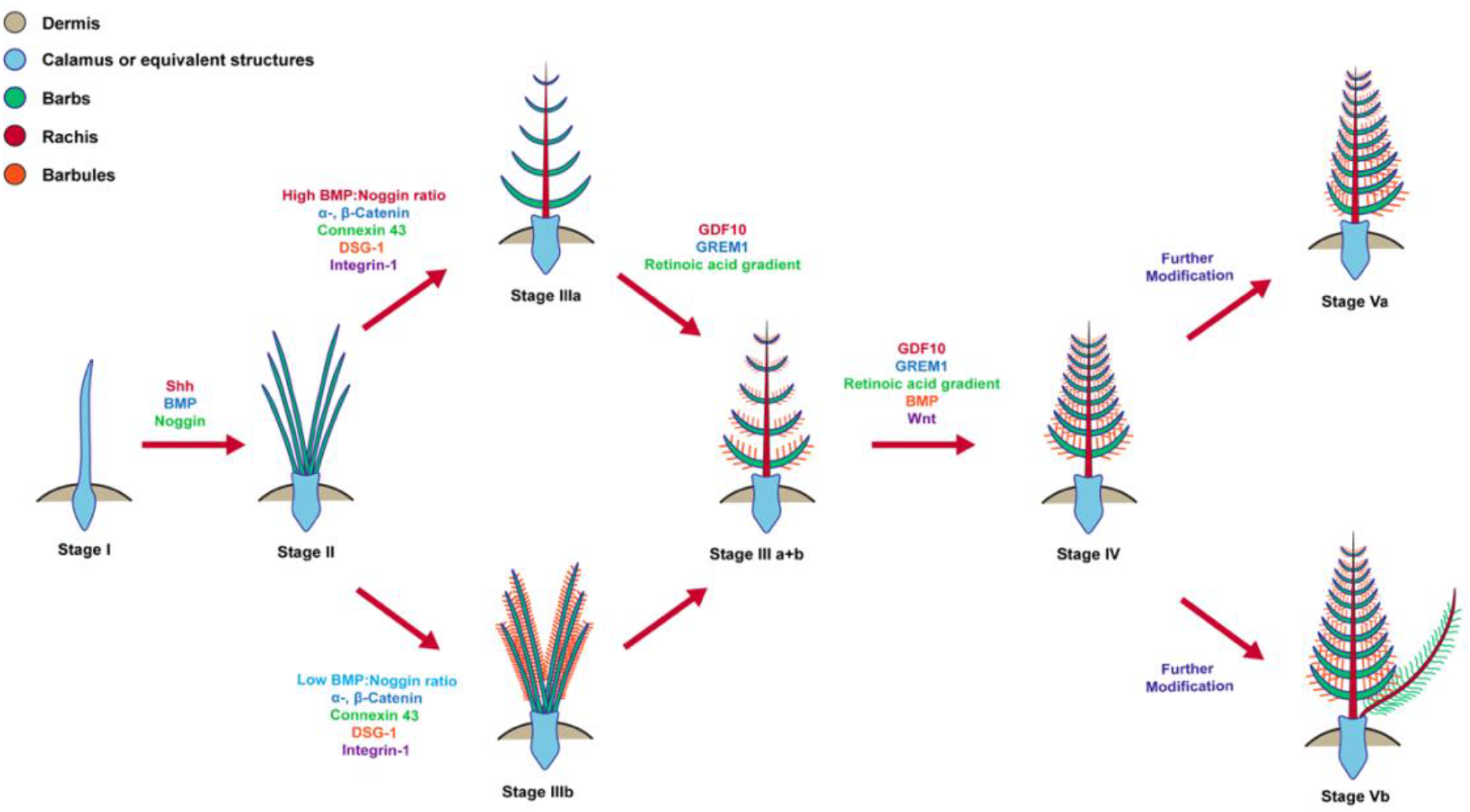
Prum and Brush model of feather morphogenesis and evolution. **Stage I** is a single undifferentiated filament. **Stage II** shows a simple branched structure with barbs. **Stage III** represents further branching in two different directions, either formation of a central rachis with barbs (**IIIa**) or numerous barbs differentiated into barbules emanating from the calamus (**IIIb**). In **Stage IV**, the barbules generate hooklets to interlock with the barbules of adjacent barbs resulting in a vaned topology. Further modification results in asymmetrically vaned flight feathers (**Stage Va**) and symmetrical vaned feathers with afterfeathers (**Stage Vb**). Other **Stage V** forms also exist but are not shown in the figure. Important molecular pathways and protein crosstalk leading to the development of various stages are added for reference. GRS Ref. 32865 is proposed as the first direct evidence of **Stage IIIb**.

The widely accepted evolution-development model of the origin of feathers proposed by Prum and Brush (2) has five stages (I-V) (Fig. 2). Stage I involves the formation of a single undifferentiated tubular filament with its base invaginating into the skin to form a cylindrical wall which later cornifies into the calamus. In this context, different stages or diverse forms of integumentary structures arise simply by tweaking various components of the signalling pathway (switching off expression, altering gradients across time or within cells, changing the ratio of competing signalling proteins etc.), i.e. by the action of a modular signalling system. Differing expression patterns of α-, β-catenin, Connexin 43, DSG-1, Integrin-1, Wnt act as molecular switches between pennaceous or plumulaceous topology (7). When both (a) and (b) take place Stage IIa+b results but their sequence is unclear (2). GDF10, GREM1 and retinoic acid gradients influence the modulation of barb-rachis angles in pennaceous feathers (8). Continued action of BMP’s in barbules leads to the formation of barbicels or hooklets, which allows the formation of closed interlocked vane (Stage IV) (9). Stage V (inclusive of true remiges, retrices, afterfeathers and many other novelties) appear on further adjustments to the Stage IV template (2).

Nested hierarchical branching is a characteristic feature of feathers. Fossil evidence of simpler, more tubular feathers in dinosaurs and the study of morphogenesis in modern bird embryos gave rise to two competing hypotheses for the evolution and development of feathers, (a) the rachis arose prior to the development of branched structures and (b) barbs arose first, and fusion of barbs leads to rachis development (3). However, evolution-development experiments clearly show that barbs and rachis formation are interconvertible during morphogenesis depending on a cascade of molecular switches (8). Down feathers are representative of Stage IIIb and the secondary development of a rachis, as evidenced by a feathered dinosaur tail from Burmese amber (10). This could have just as parsimoniously been an intermediate to Stage IIIa+b, like previously reported intermediate forms between Stage II–Stage IIIa (11).

Avian integumentary appendages have evolved in a piecemeal fashion. All nine modern feather types (simple tubular bristles, remiges, retrices, downs, contours, semiplumes, bristles, filoplumes and afterfeathers) have been reported in the fossil record (12). However, several morphologies (rachis-dominated feathers/proximally ribbon-like pennaceous feathers) have been noted in the fossil record that do not appear in modern birds (13). Evidence from quantitative morphology field analysis (QMorF), a next generation evolution-development technique, showed that tweaking protein crosstalk can generate pennaceous and plumulaceous sections even within the same feather (7). This provides strong evidence that modularity in signalling patterns and molecular crosstalk could have led to a plethora of intermediate stages yet to be discovered in the fossil record and thus makes a compelling case for the role of modularity in feather evolution. Hence, our discovery of a three-dimensionally preserved Stage IIIb feather supports the hypothesis that substantial experimentation with feather morphology was already underway in the Mesozoic. It also refutes the claim that Stage IIIb is “unlikely” to exist within the developmental roadmap of modern feathers (14).

## Materials and Methods

GRS Ref. 32865 was collected in 2019 under government permit from the Zee Phyu Gong locality (26°13’N; 96°36’E) of the Hukwang Valley of Sagaing State in northern Myanmar from rocks dated to the earliest Cenomanian stage of the Upper Cretaceous (15).The specimen GRS Ref.32865 was viewed under a Zeiss Stemi SV 11 microscope, backlit with diffused white LED light. Photographs were taken with a Nikon DSLR D850 camera attached to the microscope with the amber sample submerged in mineral oil (refractive index matching the amber) in order to minimise loss of image detail due to light refraction artefacts. The camera was controlled by digiCam Control v2.1.2.0. Images were stacked in Helicon Focus 7.6.

## Acknowledgments

We would like to thank the Peretti Museum Foundation for specimen access and study support. AR is supported by a Hong Kong PhD Fellowship (HKPF PF 16-09281). CVM. is supported by a Postgraduate Scholarship (PGS) from The University of Hong Kong. MP and TGK are supported by the RAE Improvement Fund of the Faculty of Science, The University of Hong Kong.

