## Supplementary material for "Three-dimensionally preserved ‘Stage IIIb’ fossil down feather supports developmental modularity in feather evolution": SI Appendix

Additional Notes:

(1) Collection of Fossil material and Statement of Ethics

GRS Ref. 32865 was collected in 2019 under government permit from the Zee Phyu Gong locality (26°13’N ; 96°36’E) locality of the Hukwang Valley of Sagaing State in northern Myanmar from rocks dated to the earliest Cenomanian stage of the Upper Cretaceous (1). The Hukwang Valley is ~50km west of the Tanai township (also known as Danai) which is located in neighbouring Kachin State. The mines of the Tanai area are linked with armed conflict (2, 3); to our knowledge the mines of Hukwang Valley are not (see UN Human Rights Council report (3)). Thus, we believe that GRS Ref. 32865 is ethically sourced to international standards (4).

(2) Anatomy of GRS Ref.32865 and its diagnosis as Stage IIIb

GRS Ref.32865 feather specimen measures ~1.67 mm proximodistally, with a calamus ~0.4 mm long and barbs averaging ~1.25 mm in length. It lacks a central rachis, with barbs extending radially from the calamus. The barbs appear to be flexible, as in all modern down feathers (5), allowing them to bend in a sigmoidal fashion which gives the feather a bulbous shape. Each of the barbs branch into ~50 pairs of barbules at an angle of ~28°. The angle is relatively constant as seen in modern rigid down feathers (5). This implies that barbules have a composition more similar to the rigid barbules of modern pennaceous feathers. Barbules extend to the distal tip of the barbs, a state only seen in post-natal down feathers in modern birds (5). The barbules appear to be unornamented unlike galliform, anseriform (6), or passerine (7) down feathers. The barbs do not terminate in barbicels (or hooklets) and barbules are hair-like as in Paleognathae (8). In modern birds, downy barbules tend to maintain a constant length while decreasing in basal width distally (5). However, the barbules in our specimen decrease in length near the distal end (~0.19 mm at the base and ~0.06 mm at the apex) while maintaining a fairly constant width. The calamus is stubby and seems to be encapsulated by the remnant of a decaying follicular sheath. The feather appears semi-transparent and only slightly darker than the surrounding amber when viewed using light microscopy. The type of pigmentation cannot be confirmed without destructive sampling and chemical testing. This is not possible owing to the rarity of the specimen. Hence we refrain from commenting on pigmentation but see Xing, Cockx and McKellar (9) for discussions about feather pigmentation in similar amber specimens based on observations by eye.

(3) **Burmese amber fossil record**

Fossils found in Late Cretaceous amber from Myanmar provide extensive insight into the Cretaceous tropical forests of Laurasia (10-12). Amber is fossilised tree resin (13) that was originally exuded from a tree because of mechanical damage to its bark tissue (14). Detritus and organisms can become partially embedded in an initial resin flow and subsequently encased as the flow continues (15). Subsequent burial of the resin under anaerobic conditions concentrates the resin and preserves it as amber (14). Burmese amber is believed to originate from conifers (Family: Araucareacea, phylogenetically related to the modern day *Wollemia*) (16) growing in a tropical coastal forest (10, 17) that also included angiosperms (18-22), bryophytes (23-25), pteridophytes (26, 27) as well as fungi (28-32). Animals preserved in Burmese amber include arthropods (10, 11, 33-45), nematodes (10, 11, 35, 46), molluscs (11, 47, 48), lizards (10, 49, 50) and theropod dinosaurs (particularly enantiornithine birds) (42, 45, 51-56). Burmese amber has yielded several isolated body parts referred to Enantiornithes (51, 55, 56) as well as an entire enantiornithine hatchling (53). These rare finds shed light on the soft tissue morphology and ontogenetic strategy of young enantiornithines. Fossil feathers are more common in Burmese amber but can be difficult to identify less inclusively than pennaraptoran theropods (9). Isolated feathers preserved in amber are known from the Barremian (57) of Lebanon (58), the Aptian-Albian (59) of Jordan (60) and Spain (61, 62), the Albian of France (63), the Cenomanian of Myanmar (42, 45, 54), the Turonian of New Jersey (64, 65), the Campanian of Alberta (66), and the Eocene-Miocene of the Dominican Republic (67). Only the Eocene-Miocene feather has been referred to a precise group e.g. Picidae (67), with the remainder only referable to Pennaraptora (68, 69). Most of these reports simply confirm the presence of pennaraptoran theropods in the locality, though some have provided insight into the origins of feather parasites (42, 45) and the evolution and development of the morphology of modern feathers (9, 54, 63, 66).	The oldest records of down feathers are limited to two-dimensional compression fossils from the Early Cretaceous Aptian stage of Brazil (70) and Australia (71). Burmese amber extends the record of three-dimensionally preserved fossil down feathers by at least 10 million years from the Turonian stage of the Late Cretaceous (64) or later (63, 66, 67) to the Cenomanian stage (for a complete review of the different types of feathers reported in Burmese amber see Xing, Cockx and McKellar (9).

(4) **Novelties in feathers evolved through shared molecular signalling pathways**

The widely accepted evolution-development model of the origin of feathers proposed by Prum and Brush (77) has five stages (I-V) (Fig. 2). Stage I involves the formation of a single undifferentiated tubular filament with its base invaginating into the skin to form a cylindrical wall which later cornifies into the calamus. Stage II is characterised by the division of the epidermal wall on the top of the calamus into barb ridges. Stage II feathers are therefore composed of simple barbs emanating from the calamus. The formation of barb ridges is regulated by time-dependent expression of the BMP’s and Noggin whereas barb elongation is dependent on the protein Shh (81). At this stage, one of two possible events may occur: (i) high BMP : Noggin ratio leads to helical displacement of barbs producing a rachis with unbranched barbs (Stage IIIa) or (ii) lower BMP : Noggin ratio leads to symmetrical differentiation of the barb ridges into paired barbule plates thus producing a feather without a rachis but with paired barbules (Stage IIIb) (82). Key differences in expression patterns of α-, β-catenin, Connexin 43, DSG-1, Integrin-1, Wnt and a host of other proteins also act as direct switches between either developing a central rachis and eventually a pennaceous vane or developing a plumulaceous feather topology (83).

A feather with a central rachis, barbs and symmetrically branched barbules (Stage IIIa+b) has been postulated to result from both of the above events taking place (77), but their order of occurrence and subtleties in the spatiotemporal modulation of protein-protein interactions are still unclear. Additionally, GDF10, GREM1 and retinoic acid gradients influence the modulation of barb-rachis angles in pennaceous feathers (84). Continued action of BMP’s on barbule plate cells leads to the formation of barbicels or hooklets, which allows the formation of closed vane with interlocking barbules (Stage IV) (82). Stage V (inclusive of true remiges, retrices, afterfeathers and many other novelties) appear on further modular adjustments to the Stage IV template (77).
